# Miniature spatial transcriptomics for studying parasite-endosymbiont relationships at the micro scale

**DOI:** 10.1101/2022.11.23.517653

**Authors:** Hailey Sounart, Denis Voronin, Yuvarani Masarapu, Matthew Chung, Sami Saarenpää, Elodie Ghedin, Stefania Giacomello

**Affiliations:** Department of Gene Technology, KTH Royal Institute of Technology, SciLifeLab, Stockholm, Sweden; Systems Genomics Section, Laboratory of Parasitic Diseases, National Institute of Allergy and Infectious Diseases, National Institutes of Health, Bethesda, Maryland, United States

**Author notes:** **Materials & Correspondence** Correspondence to Stefania Giacomello and Elodie Ghedin. First shared authors (Hailey Sounart, Denis Voronin).

## Abstract

Several important human infectious diseases are caused by microscale-sized parasitic nematodes like filarial worms. Filarial worms have their own spatial tissue organization and to uncover this tissue structure, we need methods that can spatially resolve these miniature specimens. Most filarial worms evolved a mutualistic association with endosymbiotic bacteria *Wolbachia*, however, the mechanisms underlying the dependency of filarial worms on the fitness of these bacteria remain unknown. As *Wolbachia* is essential for the development, reproduction, and survival of filarial worms, we focused on studying a posterior region containing reproductive tissue and developing embryos of adult female *Brugia malayi* worms. To spatially explore how *Wolbachia* interacts with the worm’s reproductive system, we performed a spatial characterization using Spatial Transcriptomics (ST) across our region of interest. We provide a proof-of-concept for miniature-ST to explore spatial gene expression patterns in small sample types, demonstrating the method’s ability to uncover nuanced tissue region expression patterns, observe the spatial localization of key *B. malayi* - *Wolbachia* pathway genes, and co-localize the *B. malayi* spatial transcriptome in *Wolbachia* tissue regions. We envision our approach to open up new scenarios for the study of infectious diseases caused by micro-scale parasitic worms.

## Introduction

The infectious diseases lymphatic filariasis (elephantiasis) and onchocerciasis (river blindness) are two debilitating and far-reaching neglected tropical diseases, affecting more than 150 million of “the poorest of the poor” worldwide^1–3^. The parasitic nematodes that cause these diseases— *Wuchereria bancrofti, Brugia malayi*, and *B. timori* cause lymphatic filariasis and *Onchocerca volvulus* causes onchocerciasis—cause severe pathologies such as blindness in onchocerciasis and lymphoedema that can progress to elephantiasis in lymphatic filariasis^3^. Mass drug administration campaigns of affordable and safe microfilaricidal drugs are used to halt parasite transmission by killing the microfilariae (mf) in the blood or the skin of the infected hosts. However, the long lifespan of the adult worms (8 years for lymphatic filariasis and 20-30 years for river blindness) requires repeated microfilaricidal treatment. Therefore, alternative therapeutic approaches that affect the adult worms are critically needed.

*Wolbachia* are common intracellular bacteria found in arthropods and filarial nematodes^4^. Filarial parasites that cause lymphatic filariasis and onchocerciasis have evolved a mutualistic association with *Wolbachia* that is essential for worm development, reproduction, and survival^5^. The endosymbiont can be eliminated from the worms by treating infected animals and humans with a small variety of antibiotics (such as doxycycline and rifampicin), which in turn results in the death of the adult worms, ultimately curing the mammalian host of the filarial infection^4,6–9^. Doxycycline treatment has demonstrated strong effects on the embryogenesis of filarial nematodes, causing apoptosis in developing embryos after 6 days of treatment *in vitro*^*10*^, and eventual embryo clearance from the uterus of females treated *in vivo* during clinical trials^3,11,12^. To study the molecular processes of the disease-causing filarial worms, *Brugia malayi* has become the model organism, as it can be cultured across all life cycle stages in the laboratory animal model *Meriones unguiculatus*^13^. A number of studies have explored at the molecular level the co-dependencies between *B. malayi* and its *Wolbachia* (*w*Bm)^14–21^. Homeostasis of the mutualistic relationship that evolved between *B. malayi* and *w*Bm requires the coordinated regulation of *B. malayi* genes. This dependency on *Wolbachia* for oogenesis and embryogenesis in female worms means that therapeutic approaches that deplete *w*Bm in the *B. malayi* worms can cause permanent sterilization of adult females. Due to the large impact *w*Bm has on the processes of oogenesis and embryogenesis within the reproductive tissue of adult female *B. malayi* worms, we need approaches that can uncover more about the molecular mechanisms occurring in this region of the worm and how such processes are spatially distributed within the different tissue structures.

Recent transcriptomic and proteomic analyses of whole *B. malayi* adult parasites have led to significant insight into the biology of the worm^22–24^. However, while comparative proteomic analysis of differences between reproductive tissue, body wall, and digestive tract regions have provided essential information for the whole tissue^22^, nuanced expression patterns in specific regions of each tissue were likely missed. Therefore, there are specific regions of the worms that are systematically underrepresented in whole-parasite omics such as the head region of the parasites that contain critical tissues at the host-parasite interface, and posterior regions where oogenesis and early embryogenesis occur. Prioritizing omic analyses of these regions will help better understand worm biology, life cycle, and transmission, as well as help gain a better handle on the interface of parasite-host interactions. Thus far, only one study showed spatial-like transcriptomic analysis of the *B. malayi* head region using a combination of highly complex methods, such as low-input tissue capture and RNA tomography, combined with light-sheet and electron microscopy^23^. Other, more standard approaches have been used to study other helminths. For example, the first cell atlas of a parasitic worm, *Schistosoma mansoni*, was made using single-cell technology^24^. However, isolating single cells for filarial nematodes such as *B. malayi* is complicated because of the large hypodermal syncytial cells spanning the entire length of the worm body, making this strategy unfeasible. Resolution of *Wolbachia* and *B. malayi* gene expression at the level of individual cells or tissue morphological regions is needed to design more effective therapeutic strategies for lymphatic filariasis. Single-nucleus RNA sequencing offers the possibility of overcoming the anatomical constraints of a single-cell based strategy, however, crucial information provided by having spatial context is lost with such a method. Thus, to resolve the spatial structures within the worm, longitudinal sections can be used with a spatial transcriptomics approach to visualize expression of *B. malayi* genes in specific tissue regions important in the *B. malayi*-*w*Bm relationship.

Spatial Transcriptomics (ST) is a high-throughput, sequencing-based exploratory method where polyadenylated transcripts are captured by spatially barcoded probes on a slide underneath a tissue section^25,26^. ST connects tissue morphology to gene expression by overlaying the spatially resolved transcriptome at 55 μm resolution on hematoxylin and eosin stained images^25,26^. This technique has been used to study a variety of tissue types and disease states across primarily human and mouse tissues^27^, as well as some plant tissues^28,29^, typically in the 1-6 mm range.

To enable spatially resolved transcriptomic methods for nematodes at the micro-scale, we focused on *B. malayi* as our model organism. We profile adult female worms that are on average 150 μm in diameter and 43-55 mm in length and focus specifically on a region containing ovary tissue, the beginning of the uterus with fertilized eggs and early embryos, the digestive tract, and the body wall. We have overcome the technical challenge of working with very small tissue sizes by developing sample preparation (fixation and embedding), cryosectioning, and tissue attachment/staining techniques for miniature Spatial Transcriptomics. We utilized such a miniature-ST method to uncover tissue-specific gene expression patterns in *B. malayi*, localize the expression of key glycolytic pathway genes, and co-localize the expression of *B. malayi* genes in tissue areas with and without *Wolbachia*. We studied gene expression patterns in both 2D and in 3D across a posterior region of the adult female worms. By developing the highly reproducible miniature-ST method, we are forging a research path to perform spatial characterization of gene expression profiles across tissues of small parasitic worms. A tissue-resolution level understanding of parasitic worm biology could help the development of more targeted therapeutic strategies.

## Results

### Miniature-ST provides reproducible capture of gene expression information

To visualize the spatial localization of *B. malayi* genes in the adult female worm, we first determined if spatial transcriptomics could be applied to an organism of this size (∼130-170 μm in diameter). We adapted the ST technology Visium Spatial Gene Expression assay (10X Genomics)^30^ for unbiased capture of gene expression information. To apply ST to this small sample type, we faced several technical challenges and developed strategies to overcome these issues in terms of sample preparation, cryosectioning, and tissue attachment to the slides. For sample preparation, we developed several steps to improve downstream cryosectioning and tissue attachment to the Visium slides. These steps involved: i) methanol fixation before embedding the tissue in the Optimal Cutting Temperature (OCT) compound to improve tissue attachment, ii) hematoxylin staining before OCT embedding to visualize the clear worms in the embedded block, and iii) dissecting out smaller, i.e. ∼5 mm, pieces greatly facilitated obtaining intact cryosections during sectioning. When cryosectioning the worm tissue, it was difficult to obtain intact, longitudinal sections from such a small specimen (less than 20 total sections per sample). We found that the sample needed to be completely flat to get usable sections through the embedded worm piece. Re-embedding of the small ∼5 mm worm specimen onto flattened OCT facilitated acquiring multiple, intact sections throughout a single worm specimen, all of which could be placed on a single Visium slide capture area, which also reduces experimental costs (**Figure 1A**). To understand if our re-embedding approach could affect the gene expression information captured, we included a sample, BM3, that was not re-embedded prior to sectioning. We found that the difference in the embedding technique did not impact the quality of the gene expression information captured as the gene expression patterns were reproducible between samples, but rather the re-embedding technique greatly aided in obtaining intact tissue sections (**Figure 1B-D**). A section thickness of 8 μm allowed us to collect essentially all the sections throughout an entire worm sample, which could then be aligned and stacked to generate a 3D model of spatial gene expression information. To address the third issue of poor tissue attachment to the Visium slides, we observed that the pre-embedding methanol fixation step, 8 μm thick cryosections, placing multiple sections on the same Visium capture area without overlapping the OCT, and modifying the H&E staining to be performed inside slide cassettes with intermittent warming, improved tissue attachment. To attain a finer image for morphological annotation, we introduced imaging with z-stacks. Overall, the implementation of these technical changes to the Visium protocol enabled us to obtain spatially resolved transcriptomic information from the worm specimens.

**Figure 1.**
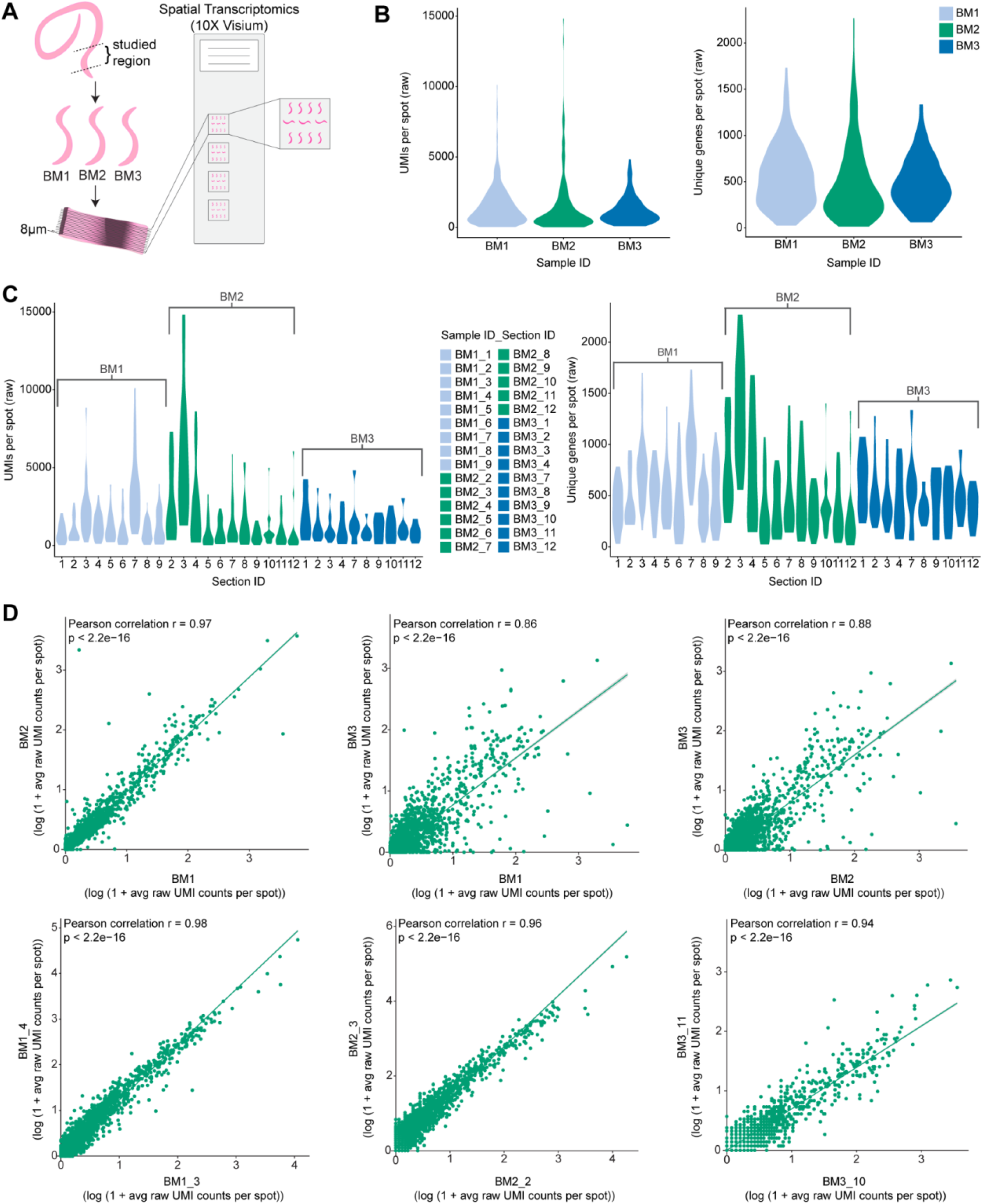
Reproducibility of miniature-ST. (A) Overview of the miniature-ST method performed in this study, where cryosections from the posterior region of adult female *Brugia malayi* worms containing ovary tissue, the beginning of the uterus with fertilized eggs and early embryos, the digestive tract, and body wall analyzed using Spatial Transcriptomics (ST, Visium). (B)Violin plots of the UMIs and unique genes per capture spot across the different worm samples used in the study. (C) Violin plots of the unique molecules (UMIs) per spot and unique genes per spot across different worm sample sections used in the study. (D) Pearson correlation of the average gene expression between worm samples (top) and sections from the same sample (bottom), p-value < 2.2e-16.

We then applied this newly developed miniature-ST technique to the tissues of selected parts of *B. malayi* adult female worms and generated a ST dataset from 3 worm samples (i.e., samples BM1, BM2, BM3) and across a total of 30 sections (**Figure 1A, Supplementary Table 1**). In total, we captured 7,724 unique *B. malayi* genes from the studied region containing ovary tissue, the beginning of the uterus with fertilized eggs and early embryos, as well as part of the digestive tract and body wall. These represent 66% of the 11,777 *B. malayi* genes in the current genome annotation (WormBase: WBPS14), across 547 spots with an average of ∼1,457 unique molecules (UMIs) per spot and ∼529 unique genes per capture spot. The number of genes we captured is in the same range as the 8,000-10,000 genes (70-90% of genome) per sequence library captured by a previous bulk RNAseq study across whole worms and different lifecycle stages^31^. We observed similar unique gene and unique molecule distributions per spot across the different samples (**Figure 1B**) and across sections from the same sample (**Figure 1C**). In addition, different samples (r = 0.86-0.97, p < 2.2e-16) and different sections from the same sample (r =0.87-0.98, p < 2.2e-16) had a high correlation in their average gene expression (**Figure 1D**). Such results demonstrate that spatial transcriptomic information was reproducibly captured across samples and sample sections with our miniature-ST method.

### Spatially distinct gene expression separation of digestive tract, body wall, and multiple reproductive tissue regions

Given the high reproducibility of the approach, we then performed unsupervised clustering analysis of the miniature-ST data and identified four distinct clusters (**Figure 2A-B**). Differential expression (DE) analysis identified marker genes for each cluster, with 36 DE genes in cluster 1, 82 DE genes in cluster 2, 58 DE genes in cluster 3, and 31 DE genes in cluster 4 that significantly changed their expression in each cluster as compared to the other 3 clusters (**Supplementary Table 2A**). The significantly upregulated marker genes (positive logFC, p<0.05) in each cluster were annotated using a previous proteomics study^22^, which identified each cluster as specifically enriched in a set of highly expressed tissue-specific markers (Methods). Thus, we considered each cluster as representative of the tissue type for which the cluster contained highly expressed tissue specific markers: digestive tract for cluster 1, reproductive tissue for clusters 2 and 4, and body wall for cluster 3 (**Figure 2C, Supplementary Table 2B-F**). Of note, cluster 3 was a mixed cluster with higher enrichment in body wall marker genes (9.3%), but also contained markers for the reproductive tract (4.7%) (**Figure 2C, Supplementary Table 2B-C**). When looking at the spatial distribution of cluster 3 on the tissue sections (**Figure 2B**), we observed that the spots localized to both tissue types identified at the gene expression level: the body wall (hypodermal chord, muscles) found towards the exterior part of the worms, and the reproductive tissue located inside the worms. Thus, with a resolution of 55 μm we observed at both the spatial spot localization level and at the gene expression level a tissue-specific separation by clustering.

**Figure 2.**
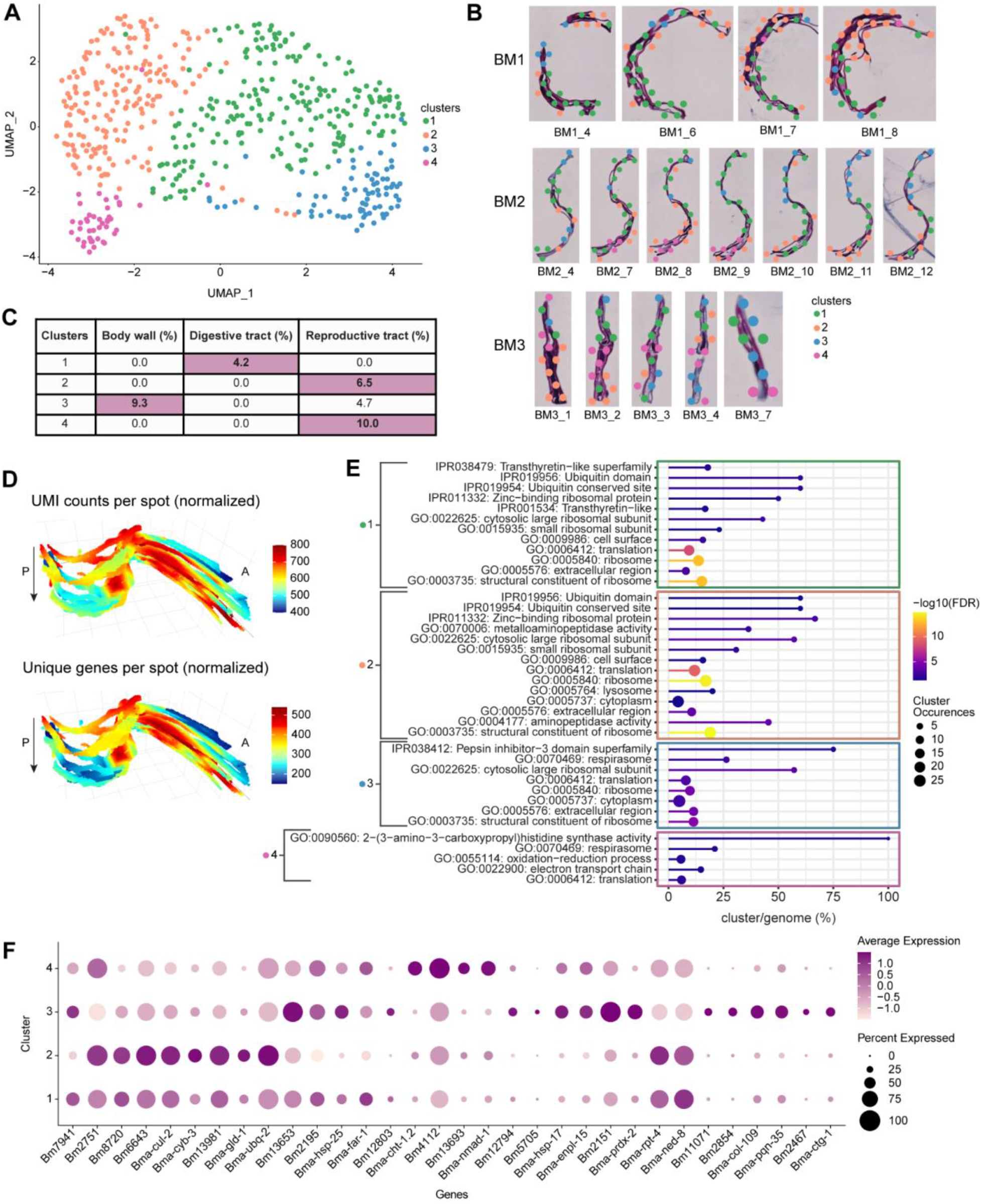
Tissue region specific spatial gene expression patterns. (A) UMAP showing the four clusters of *Brugia malayi* transcriptome data. (B) Spatial distribution of the four clusters on tissue sections from *B. malayi* samples. Magnification 20x. (C) Percent of differential expressed marker genes per cluster enriched for specific tissue regions. (D) 3D model of the UMIs and unique genes per spot across worm BM2. Magnification 20x. A: Anterior, P: Posterior, arrow indicates first to last section through the worm. (E) Significantly overrepresented functional terms for each cluster (FDR<0.05) (F) Dotplot depicting expression of *B. malayi* cluster marker genes across the different clusters.

To study how gene expression patterns localize to different morphological structures and to see if there are specific patterns that arise across the entire worm region, we explored different cluster marker gene spatial distributions in 3D by compiling all the consecutive sections from worm sample BM2 (**Figure 2D**). By visualizing the expression patterns of marker genes for the digestive tract, the reproductive tract, or the body wall, we could localize the corresponding tissue region in the 3D reconstruction (**Supplementary Figure 1**). Furthermore, we performed fixed term enrichment analysis to provide an overview of the genes and processes enriched in each cluster (**Figure 2E, Supplementary Table 3A-B**). Cluster 1 contained marker genes associated with the cell surface and interactions between cells and the environment. The most significant DE marker genes for cluster 1 were ones with ribosomal function and its components, ubiquitin functions, and Transthyretin-like family proteins (p<0.001) (**Figure 2E, Supplementary Table 3A-B**). Proteins annotated as Transthyretin-like proteins are associated with the cell surface in *B. malayi* and Transthyretin is a protein that transports the thyroid hormone thyroxine and Vitamin A (retinol). In terms of interactions between cells and the environment, we observed a positive expression of genes encoding proteins involved in ion binding activity, ion transmembrane transporter activity, sterol-binding and transfer activity, and ShKT-domain containing proteins (**Figure 2E, Supplementary Table 3A-B**). The largest family of proteins containing ShKT-line domain is found in worms *Caenorhabditis elegans, C. briggsae, B. malayi, B. pahangi, Ancylostoma ceylanicum, S. mansoni*, and *Toxocara canis*^32,33^. ShKT-domain containing proteins from parasitic nematodes were recently shown to possess immunomodulatory activity via the blockage of voltage-gated potassium channels on human effector memory T cells^32^. It was also shown that some ShKT-like domain containing proteins were highly expressed along the digestive tract in *C. elegans* adult worms^34^. Additionally, the expression of a marker for the digestive tract (cluster 1), Bm7941, changed throughout the entire 3D worm sample. Specifically, when moving through the 3D model of the worm sample, the elevated expression of Bm7941 shifted from the anterior side to the posterior side, further supporting cluster 1 as localizing to the digestive tract (**Supplementary Figure 1, Supplementary Table 2D**).

According to a proteomic analysis, three genes (Bm2751, Bm8720, Bm6643) upregulated in cluster 2 are markers for reproductive tissue (**Figure 2F, Supplementary Table 2A**,**E**). The expression of these genes in cluster 2 was elevated in the middle, closer to the posterior region, and then shifted towards the anterior part of the worm through the 3D model (**Supplementary Figure 1, Supplementary Table 2E**). Cluster 2 covers a substantial internal part of the worm and, according to highly expressed reproductive tissue markers, represents a tissue region containing the ovaries and the uterus. We observed that more than 13% of all up-regulated genes in this cluster are involved in processes associated with oogenesis and early embryogenesis (**Supplementary Table 2A**). According to their *C. elegans* orthologs, these genes are part of the following processes: meiotic chromosome segregation and organization (Bma-cul-2, Bma-cyb-3), oocyte maturation (Bma-cyb-3), gamete generation (Bm13981), positive regulation of female gonad development (Bma-gld-1), and polarity specification of the anterior-posterior axis (Bma-ubq-2) (**Figure 2F, Supplementary Table 2A**). This pattern confirms that cluster 2 truly represents the reproductive tissue of worms. Down-regulated genes in cluster 2 consist of heat-shock proteins (Bm13653, Bm2195, Bma-hsp-25) and lipid binding and lipid droplet disassembly processes (Bma-far-1, Bm12803) (**Figure 2F, Supplementary Table 2A**). During oogenesis, lipids are stored for later use in rapid early embryogenesis. Therefore, the down regulation of genes involved in the degradation of stored lipids indicates that cluster 2 is closer to the ovaries than the uterus region of the reproductive tract.

Clusters 3 and 4 also contained markers for the reproductive tract but appeared to localize to distinct reproductive tract tissue regions. Markers Bma-cht-1.2 and Bm4112 for cluster 4 partially overlapped the cluster 2 reproductive tract markers expression pattern; clusters 2 and 4 were located closer to one another both in UMAP space and spatially on the tissue sections (**Figure 2A-B**). However, cluster 4 markers could represent a different reproductive tract region than cluster 2 (**Figure 2F, Supplementary 2A**,**F**). After oocytes are fertilized in the adult female worm reproductive tract, embryos build a shell mainly composed of chitin and develop into microfilaria (a pre-larval worm stage) within this chitin shell. Bma-cht-1.2 is involved in chitin binding activity and thus the region where Bma-cht-1.2 showed elevated expression may represent the viaduct region of the reproductive tract, which connects the ovaries to the uterus; it is where fertilization and very early formation of the embryos occur (**Supplementary Figure 1, Supplementary Table 2F**). In addition, cluster 4 displayed significant enrichment of genes encoding proteins with respiration activity, including the electron transport chain, respirasome and oxidative-reduction process, histone (H4), and histone binding proteins synthesis (**Figure 2E, Supplementary Table 3A-B**). Histones and histone-binding proteins, such as those encoded by Bm4112 and Bm13693, are important for developing embryos in the reproductive tract, which corresponds to the region used in this study. The histone H4 (Bm4112) upregulated in cluster 4 (p=0.0004) is an ortholog of *C. elegans* his-10, his-31, and his-64 (**Figure 2F, Supplementary Table 2F**). In *C. elegans*, his-10 is responsible for chromatin formation and involved in the defense response to Gram-negative bacteria and the innate immune response. In cluster 4, we also observed higher expression of genes essential in oogenesis, such as Bma-nmad-1, an ortholog of *C. elegans* nmad-1 (**Figure 2F, Supplementary Table 2A**). In *C. elegans*, nmad-1 is involved in meiotic chromosome condensation, positive regulation of organelle organization, and positive regulation of oviposition. Due to the upregulation of genes involved in oogenesis and fast-dividing embryos in the early steps of embryogenesis, we conclude that cluster 4 represents the region of the uterus with fast dividing cells of early embryos and this region is adjacent via the viaduct to the ovaries (cluster 2).

Cluster 2 likely represents the ovarian part of the reproductive tract, cluster 4 the viaduct and uterus portion of the reproductive tract, while cluster 3 contains markers (Bm12794, Bm5705) that represent a third, distinct region of the reproductive tract (**Figure 2F, Supplementary Table 2C**). Bm12794 and Bm5705 show elevated expression in the posterior part of the worm, a largely distinct pattern from the central, ovarian (cluster 2) and viaduct (cluster 4) reproductive tract marker expression patterns (**Supplementary Figure 1**). The cluster 3 reproductive tract region could correspond to the uterus. Like cluster 1 (digestive tract), cluster 3 includes a substantial set of upregulated heat-shock proteins (Bm13653, Bma-hsp-17, Bma-hsp-25) (**Figure 2F, Supplementary Table 2A**). Activation of heat shock proteins in the cluster correlated with an upregulation of unfolded protein binding activity (Bma-enpl-1) and peroxidase activity (Bm2151, Bma-prdx-2). However, proteasome-mediated protein catabolic activity (genes: Bma-rpt-4, Bma-cul-2) is downregulated in this cluster. Finally, cluster 3 showed a downregulation of Bma-ned-8 (**Figure 2F, Supplementary Table 2A**). Likely active stress-related processes found in cluster 3 result in the downregulation of Bma-ned-8. In *C. elegans*, ned-8 is involved in the regulation of the apoptotic process through signal transduction by a p53 class mediator. As cluster 3 also has markers of the reproductive tract, we assumed that ned-8 is more involved in this tissue. It was shown that elimination of *w*Bm from *B. malayi* initiates programmed cell death in the germline in ovaries and in the embryos in the uterus^10^. The induction of apoptosis was determined through the increase of expression of the cell death protein-3 (*ced*-3) gene, and the increase in the amount of inactive and active (cleaved) CED-3 protein forms in antibiotic treated *B. malayi* worms as compared to untreated (control) worms^10^. In addition, cluster 3 had 2x the number of markers for the body wall (Bm11071, Bm2854, Bma-hsp-17) than markers for the reproductive tract (**Figure 2C,F, Supplementary Table 2B-C**), where the body wall consists of muscle, cuticule, hypodermal cells, and nerve cells. These body wall markers (Bm11071, Bm2854, Bma-hsp-17) showed elevated expression in the outer areas of the worm in the first and last sections of the stack (**Supplementary Figure 1**), an expected expression pattern considering the first and last sections will likely contain the outermost portions of body wall when collecting longitudinal sections. Furthermore, Bm11071 and Bm2854 encode predicted cuticle structural constituent proteins as the cuticle forms the outermost region of the body wall, their expression pattern further supports cluster 3 as partially localizing to the body wall region of the worm. In addition to the overexpression of genes that are structural constituents of the cuticle (Bm2854, Bma-col-109, Bm11071), Bma-pqn-35, a gene ortholog of a *C. elegans* gene expressed in muscle cells, was also overexpressed in cluster 3 (**Figure 2F, Supplementary Table 2A**). The upregulation of genes encoding cuticle and muscle-related genes further supports the partial localization of cluster 3 to the body wall tissue region.

We observed an elevation of expression of Bm2467 and Bma-ctg-1 genes in cluster 3, indicating the increase of fatty acid metabolic processes in the body wall region of the parasite as Bm2467 is predicted to enable 3-hydroxyacyl-CoA dehydrogenase activity and enoyl-CoA hydratase activity, and Bma-ctg-1 is a lipid binding protein (**Figure 2F, Supplementary Table 2A**). As the body wall consists of high metabolic tissues, such as muscle and hypodermal cells, the higher activity of metabolic processes is expected in these regions. Interestingly, metabolic processes for lipids/fatty acids turnover were the most significant. Recently, it was shown that *Wolbachia* residing in the hypodermal cells of the parasites induces and uses the glycolytic pathways^19,35^.

We hypothesize that fatty acid metabolism may also be involved in providing carbohydrates to symbiotic bacteria. Overall, the clustering analysis facilitated an exploration of nuanced spatial gene expression patterns, as we could spatially pinpoint the expression of genes and pathways that were differentially regulated in the digestive tract, the body wall, and multiple reproductive tract tissue regions.

### Spatial localization of key *B. malayi* glycolytic enzyme genes

We next focused specifically on the spatial expression patterns of key glycolytic enzyme-related genes that are important in the *B. malayi* - *Wolbachia* mutualistic relationship. Pyruvate is one of the most essential metabolites for prokaryotic cells. *Wolbachia* has the full complement of genes that can use pyruvate for gluconeogenesis and for energy metabolism via the tricarboxylic acid cycle (TCA cycle)^35^. However, *Wolbachia* is missing some key enzymes that are needed for making pyruvate^19^. In previous studies, we showed that glycolysis and other pathways that produce pyruvate in filarial worms (such as *B. malayi*) provide pyruvate to *Wolbachia*^19,35^.

Glycolytic enzymes were shown to play an important role in maintaining the mutualistic association between *w*Bm and *Brugia* worms. We analyzed the genes involved in *B. malayi* pyruvate metabolism between clusters and between different tissues of the worm. We hypothesized that we could define the source and location of the pyruvate used by *Wolbachia*. We looked specifically for genes associated with glycolysis (the pathway that produces pyruvate), gluconeogenesis (the pathway where pyruvate is used to synthesize glucose, and precursors for nucleotide biosynthesis), lactate dehydrogenase (where lactate is converted to pyruvate and the reverse), and enzymes that convert cysteine amino acids to pyruvate (**Supplementary Table 4**).

Although many glycolysis genes were captured by our miniature-ST method and can be explored in our shiny app (https://giacomellolabst.shinyapps.io/brugiast-shiny/), we focused on three genes (Bma-aldo-1, Bm5699, Bma-ldh-1) that were found among our cluster marker DE genes (**Figure 3A, Supplementary Table 2A**). Bma-aldo-1(aldolase-1) and Bm5699 (glyceraldehyde 3-phosphate dehydrogenase) encode an enzyme for glycolysis, and showed increased expression in the digestive tract (cluster 1) and body wall (cluster 3) clusters, but were downregulated in the reproductive tract clusters (clusters 2 and 4) (**Figure 3A-C, Supplementary Table 2A**). *w*Bm are located in the hypodermal cells (part of the body wall) and in the ovaries, oocytes and developing embryos within the uterus. However, it is most abundant in the lateral chord (part of the body wall) as compared to the oocytes and embryo, where few *w*Bm are found. We suspect that presence of *Wolbachia* in the body wall, where bacterial load is at its highest, could induce the expression of glycolytic enzymes, resulting in the production of pyruvate thus ensuring bacterial survival; this is not the case in reproductive tissue^16,19,35^. The third gene, Bma-ldh-1 (lactate dehydrogenase or LDH) showed elevated expression in cluster 2 (reproductive tract) and was downregulated in cluster 3 (reproductive tract and body wall) (**Figure 3A,D, Supplementary Table 2A**). LDH converts lactate to pyruvate and pyruvate to lactate, which may be a reason for the differential expression of this enzyme between clusters 1 and 3.

**Figure 3.**
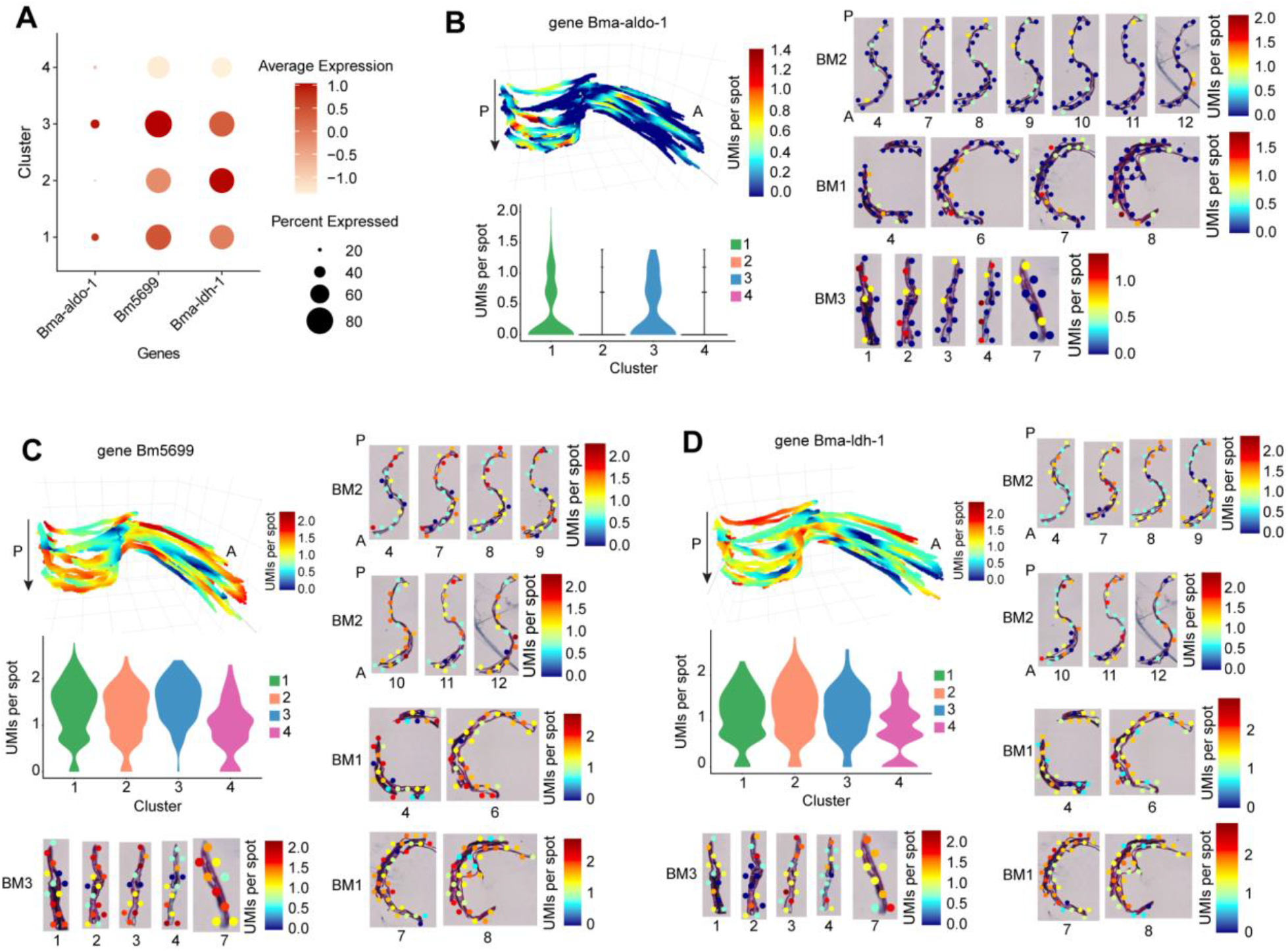
*B. malayi* glycolytic enzyme genes spatial distribution. (A) Dotplot depicting expression of glycolysis genes across the different clusters. (B-D) 3D model of gene expression in sample BM2, 2D gene expression across select clusters from all 3 samples, and gene expression distribution in a violin plot across the clusters for glycolysis genes Bma-aldo-1 (B), Bm5699 (C), and Bma-ldh-1 (D). Magnification 20x. A: Anterior, P: Posterior, arrow indicates first to last section through the worm.

### Co-localization of *B. malayi* genes in *Wolbachia*+ compared to *Wolbachia*-regions of the tissue

When capturing polyadenylated transcripts using the miniature-ST method, we detected unspecific capture of *Wolbachia* tRNA and rRNA and inferred that this signal would be indicative of *w*Bm localization. We ran our miniature-ST library reads against the *Wolbachia* tRNA and rRNA gene sequences, as those are typically the most highly abundant transcripts in an organism. We detected at least 1 UMI from 33 tRNA/rRNA *Wolbachia* genes across 206 spots (39.5% of spots) in the worm samples (**Figure 4A**). We colocalized *B. malayi* gene expression in relation to *Wolbachia* by determining the *B. malayi* DE genes between spots containing *Wolbachia* (*Wolbachia*+) and spots lacking *Wolbachia* (*Wolbachia*-) in worms and identified 65 DE genes (p<0.05) with 48 downregulated genes and 17 upregulated genes (**Figure 4B, Supplementary Table 5**). The presence of *Wolbachia* in the tissue is associated with a significant downregulation in expression of *B. malayi* genes that are predicted to encode proteins with functions that include protease and peptidase activity, endopeptidase activity, and proteosome-mediated protein catabolic process. This results in the potential reduction of proteolysis in *Wolbachia*+ tissue of female worms. We hypothesized that the inhibition of proteolysis in *Wolbachia*+ tissue protects bacterial proteins, including those that are involved in the interaction with the host.

**Figure 4.**
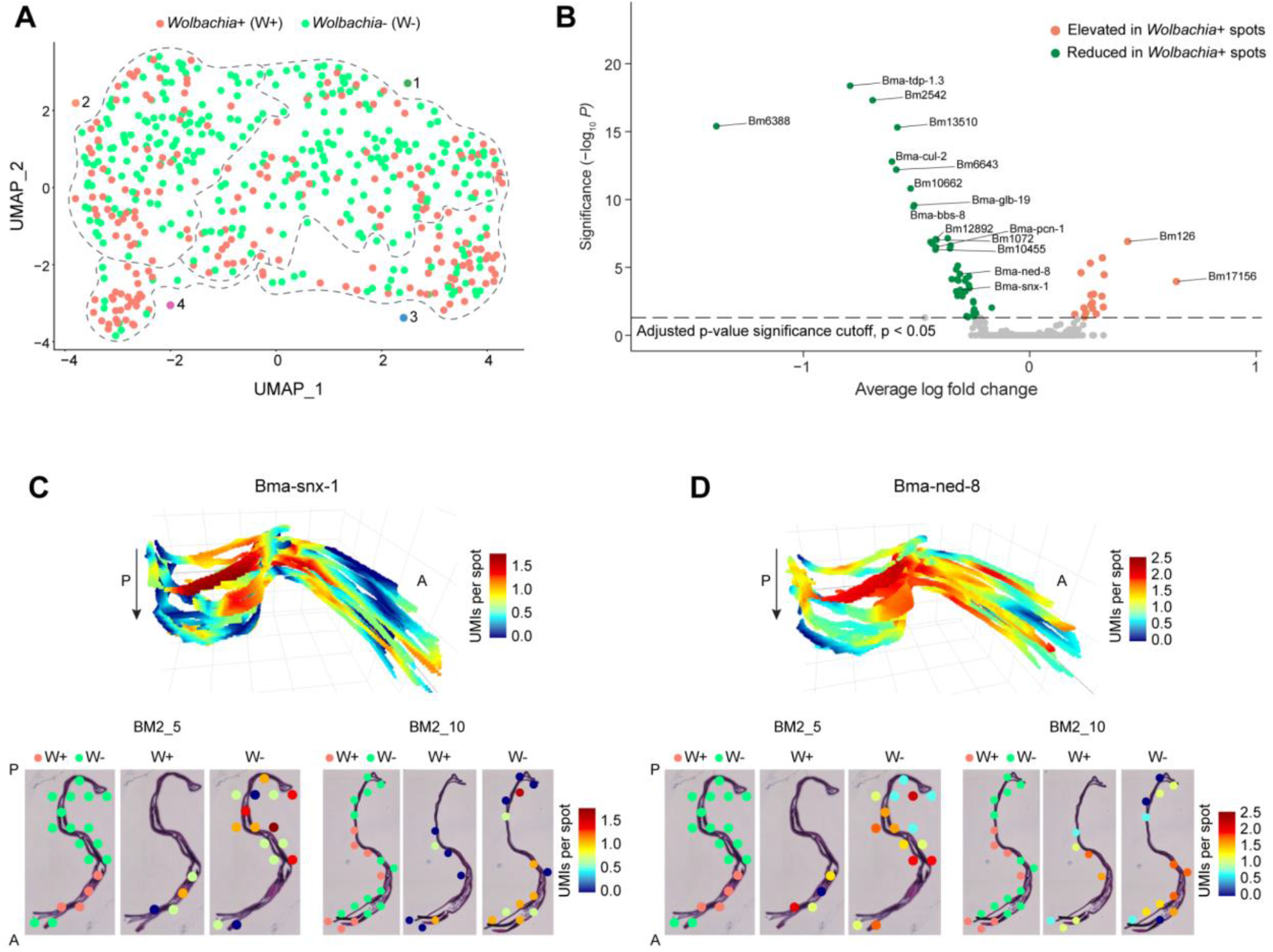
Co-localization of *B. malayi* genes in *Wolbachia*+ versus *Wolbachia*-spots. (A) UMAP showing spots with presence (*Wolbachia*+/W+) or absence (*Wolbachia*-/W-) of *Wolbachia*. Dotted lines outline the different clusters from Figure 2A. (B) Volcano plot showing the differentially expressed genes in *Wolbachia*+ versus *Wolbachia*-spots. (C-D) 3D model and 2D images of gene expression in sample BM2 for two differentially expressed genes, Bma-snx-1 and Bma-ned-8 (D), between *Wolbachia*+ (W+) and *Wolbachia*- (W-) spots. Magnification 20x. A: Anterior, P: Posterior, arrow indicates first to last section through the worm.

Across the downregulated genes in *Wolbachia*+ spots, two genes caught our attention: Bma-snx-1 and Bma-ned-8 (**Figure 4B-D, Supplementary Table 5**). Bma-snx-1 is an ortholog of *C. elegans* snx-1, and it is predicted to enable phosphatidylinositol binding activity (SNX). SNX proteins promote phagosome-lysosome fusion and are involved in apoptosis and autophagy-mediated elimination of apoptotic cells^36,37^. The inhibition of SNX proteins in *Wolbachia*+ tissue indicates that *Wolbachia* could be suppressing the late steps of autophagy and endosomal/phagosomal degradation by blocking their fusion with lysosomes. *Wolbachia* itself is surrounded by a host-derived vacuole and can be recognized as a phagosome in the cytoplasm of host cells^38^. Therefore, SNX proteins could play a role in protecting *Wolbachia* in the cytoplasm of eukaryotic cells. Bma-ned-8, an ortholog of *C. elegans* ned-8, contains ubiquitin-like domains and is also involved in regulation of the apoptotic process during embryogenesis of *C. elegans* worms^39^. As these genes are downregulated in tissue with *Wolbachia*, we hypothesized that the presence of *Wolbachia* can decrease apoptotic processes in developing embryos and ned-8 plays a key role in this regulation (**Figure 4B-D, Supplementary Table 5**). It is known that the elimination of *Wolbachia* increases intensive apoptosis in germ cells and in developing embryos of antibiotic-treated females (7 days treatment)^10^. However, elimination of the bacteria reduces support of worm biological processes and, consequently, could induce programmed cell death in the reproductive system of the worms. It is also possible that the elimination of *Wolbachia* has a direct effect on the expression of genes that regulate apoptotic processes in *B. malayi* worms.

## Discussion

In this work, we present miniature-ST, a method to analyze the spatial transcriptome of samples on the micrometer scale. We modified embedding, cryosectioning, fixation and staining steps to enable the analysis of the spatial structures composing small pathogens, such as parasitic filarial worms, that cause a variety of infectious diseases. Our spatially-resolved characterization of filarial parasitic nematode *Brugia malayi*’s transcriptome in a posterior region of the worm, in addition to the spatial capture of its endosymbiont *Wolbachia*, unveiled 2D and 3D tissue-specific gene expression patterns and the co-localization of *B. malayi* genes to specific tissue regions containing *Wolbachia*.

We applied our miniature-ST approach to visualize the spatial localization of *B. malayi* genes along a region in the adult female worms (contains ovary tissue, the beginning of the uterus with fertilized eggs and early embryos, part of the digestive tract, and body wall) in a series of 30 cryo-sections across three different worms. We captured 66% of the genes in the annotated *B. malayi* genome with similar expression profiles across samples and sections, demonstrating the ability of the miniature-ST method to reproducibly capture a significant portion of the worm’s transcriptome with spatial resolution. We identified an enriched set of tissue specific markers – annotated according to a previous proteomics study^22^ – in each cluster, indicating each cluster represented a distinct tissue type: digestive tract for cluster 1, reproductive tissue for clusters 2 and 4, and body wall for cluster 3. By constructing a 3D model of all consecutive sections throughout an entire worm sample, we observed that these tissue-specific gene expression patterns also shifted in the third dimension/z-plane, from anterior to posterior, through the worm region. Furthermore, fixed term enrichment analysis revealed genes and processes enriched in each cluster/tissue. Specifically, the analysis showed the expression of genes associated with the cell surface, cuticle, and interactions between cells and the environment in the cluster 1 for the body wall; cluster 2 for the reproductive tissue contains more than 13% of all up-regulated genes that are involved in processes associated with oogenesis and early embryogenesis. Expression of marker genes for both clusters also shifted through the z-plane of the 3D model of the worm, further supporting the cluster’s tissue specific localization. In addition, we showed the spatial localization of key *B. malayi* glycolytic enzyme genes that are essential for *Wolbachia*-worms symbiosis^19,35^. For example, two glycolytic enzyme genes (Bma-aldo-1(aldolase-1) and Bm5699 (glyceraldehyde 3-phosphate dehydrogenase)) were observed to be upregulated in *Wolbachia* abundant tissue regions and downregulated in more *Wolbachia* deficient tissue regions, which led us to hypothesize that *Wolbachia* may induce the expression of glycolytic enzymes.

Increased expression of glycolytic enzymes results in the production of pyruvate, of which *Wolbachia* is dependent on *B. malayi* for, and thus could be a mechanism to ensure bacterial survival. By spatially localizing key *B. malayi* - *Wolbachia* relationship genes, we provide a new spatial, three dimensional context to the *B. malayi* - *w*Bm relationship. Miniature-ST facilitates the separation of different tissue regions, opening up the possibility of exploring genes and pathways of interest within the distinct spatial structures and across the three dimensional reconstruction of these parasitic worms.

To further explore the *B. malayi* - *w*Bm relationship, we identified regions of the tissue containing *Wolbachia* (*Wolbachia*+ spots), leveraging unspecific capture of *Wolbachia* transcripts by poly-d(T) capture, and explored how *B. malayi* genes are modulated in different tissue regions in the presence and absence of *Wolbachia*. Our co-localization analysis unveiled a potential reduction of proteolysis in the *Wolbachia*+ tissue of the female worms, protecting bacterial proteins from degradation and thus maintaining the bacterial interaction with the *B. malayi* host. Moreover, our data illuminated new insights into understanding the role of autophagy in regulation of *Wolbachia*-host symbiosis^38^. Autophagy is a key host intracellular defense mechanism that regulates the bacterial population abundance inside host cells. Co-localization analysis of *B. malayi* genes suggests that *Wolbachia* could be suppressing the late steps of autophagy and endosomal/phagosomal degradation by blocking phagosomal fusion with lysosomes. Specifically, SNX proteins (promote phagosomal fusion with lysosomes, autophagy, and apoptosis) expression was downregulated in *Wolbachia*+ spots, and thus could play a role in protecting *Wolbachia* in the cytoplasm of eukaryotic cells. Even though our capture of *Wolbachia* was unspecific, we could spatially distinguish worm tissue regions containing *Wolbachia* and elucidate the regulation of *B. malayi* genes in those areas with miniature-ST. Furthermore, our 3D model further refined the spatial distribution of the genes involved in these molecular processes, providing new, multidimensional spatial context to their expression in relation to *Wolbachia*. Future studies exploring the spatial co-localization of *B. malayi* gene expression in relation to *Wolbachia* could further address the symbiotic relationship of these two organisms. For example, future experiments will involve studying antibiotic treated worms to understand how antibiotic-induced *Wolbachia* clearance impacts gene expression of the worms. An ideal next step would be to directly identify *Wolbachia* genes in the worms to facilitate more in-depth co-localization studies, especially in response to antibiotic treatment.

The miniature-ST method could be extended to the full *B. malayi* adult worm or to other developmental stages. Additional studies could include parasitic worms within human tissue, providing a spatial exploration of the parasite actively infecting host tissue and opening up the possibility of studying multilevel organism co-localization (*Wolbachia* endosymbionts inside the parasite and the parasite inside human host tissue). Limitations of the study include a spatial resolution of 55 µm, which does not yet allow single cell resolution. Imaging based spatial approaches offer the possibility of high, even subcellular, resolution; however, such methods focus on a specific set of target genes, making it difficult to design gene probes for species with a poorly annotated reference. ST, as a sequencing-based approach, is highly suitable for species that are not well annotated. Although working with small samples poses technical challenges, an advantage is the possibility of collecting all sections through the entire worm to construct a 3D model of the gene expression patterns throughout the entire worm sample. Additionally, 3D reconstruction of spatial gene expression throughout the length of an entire worm, all tissue types included, would provide a wealth of information to explore. A 3D model not only confirms the localization of certain expression patterns to specific morphological regions, but also can provide new insights based on how these patterns shift throughout the entire specimen. Constructing a tissue atlas from small-scaled tissue is also economically advantageous, since all tissue sections can be collected for one sample on a single Visium capture area. The spatial gene expression information we collected across all sections from *B. malayi* is presented in a shiny app (https://giacomellolabst.shinyapps.io/brugiast-shiny/), offering the possibility for further exploration.

In conclusion, we present miniature-ST, a method to unlock the potential for exploratory studies of small infectious pathogens at a spatial scale. We provide the spatial characterization of *B. malayi* gene expression in a region of adult female worms, an important step in being able to spatially resolve micro-scale disease-causing pathogens that have their own spatial tissue structure. We not only spatially resolved the parasitic worm *B. malayi*, but also its endosymbiotic bacteria, *Wolbachia*, opening up the possibility of exploring how *B. malayi* gene expression is spatially co-localized to *Wolbachia*. By spatially capturing the *B. malayi* transcriptome, this study forges a new path for studies that aim to spatially resolve gene expression information in other parasitic worms responsible for human infectious diseases. We envision our miniature-ST approach can be readily applied to other small pathogenic worms, providing new insights into the spatial organization of gene expression in these instigators of a multitude of infectious diseases.

## Methods

### Ethics statement

All animal work conducted by the NIH/NIAID Filariasis Research Reagent Resource Center (FR3) followed the national and international guidelines outlined by the National Institutes of Health Office of Laboratory Animal Welfare and was approved by the University of Georgia Athens.

### Parasite material and treatment

*B. malayi* parasites recovered from the peritoneal cavity of infected gerbils (*Meriones unguiculatus*) were obtained from FR3, University of Georgia, Athens. Adult *B. malayi* female worms were placed in 7 ml of complete culture medium (RPMI-1640 supplemented with 10% FBS, 100 U/mL penicillin, 100 mg/mL streptomycin, 2 mM L-glutamine), 2 worms per well in 6-well plate, and incubated at 37°C under 5% CO2 conditions.

Worms were incubated in the media for 2-3 days to assure they were alive, then worms were washed 3 times in PBS and fixed in 100% cold methanol (−20°C) for 5 min. Fixed samples were briefly stained with hematoxylin. This step is required to facilitate the visualization of the worms in an OCT block in future steps. A region of interest was then cut (**Supplementary Figure 2**) and placed in a vinyl specimen mold followed by the addition of OCT compound, and immediately frozen. Blocks were stored at −80°C.

### Total RNA extraction - RNA quality

To verify the RNA quality of the *B. malayi* samples, we extracted RNA from a portion of the worm. Methanol fixed, OCT-embedded *B. malayi* adult female tissue (see section “Parasite material and treatment”) was cryosectioned longitudinally to 8 µm thickness with a CryoStar NX70 cryostat (ThermoFisher). All sections (∼11-16 sections per block) throughout 2-3 embedded worm pieces were collected in Lysing Matrix D tubes (MP Biomedicals, 2mL, Cat Ref. 6913100). The extraction was done with replicates of the originally embedded blocks and re-embedded blocks where the worm piece was re-embedded onto flattened OCT on a cryo chuck from the same batch as those used for Spatial Transcriptomics. The sections in Lysing Matrix D tubes were stored at −80°C overnight. Total RNA was extracted using the RNeasy Plus Mini Kit (Qiagen, Cat No. 74136) with the modifications that 350 µl Buffer RLT Plus + β-mercaptoethanol was added to each Lysing D matrix tube (5 µl of β-mercaptoethanol (2-Mercaptoethanol, Sigma-Aldrich, M6250), added to 500µl of Buffer RLT Plus (from kit) per sample). The samples were run in a FastPrep-24 5G (MP Biomedicals) for 40 seconds at 6.0 m/sec, centrifuged for 5 min at 12000 rpm. The water phase was collected and transferred to a gDNA Eliminator spin column (provided in the kit) that was sealed with parafilm and centrifuged for 30 s at 10000 rpm. The flow-through containing RNA and proteins was transferred to an eppendorf tube (provided by kit); the beads were added to the same gDNA Eliminator column, and the column sealed with parafilm and centrifuged for 30 s at 10000 rpm. The flow-through containing RNA and proteins was then transferred to the previous aliquot in the eppendorf tube and 250 µl 96-100% ethanol (Ethanol 96%, VWR, #20823.290 and Ethanol absolute, VWR, #20816.298) was added. The manufacturer’s instructions were then followed from the RNeasy Plus Mini Handbook (09/2020) protocol for tissue samples from step 6, including the optional step 10, through step 11 and the RNA was eluted in 10 µl RNase free water (provided by kit). The concentration of extracted total RNA was determined with the RNA HS Qubit assay (Invitrogen by Thermo Fisher Scientific, REF Q32852) following the manufacturer’s instructions. Total RNA was diluted, when needed, to between 2-5ng and RIN values determined using the Agilent RNA 6000 Pico Kit following the manufacturer’s instructions. The originally embedded samples showed an RNA quality of 6.2-7.9 RIN and the re-embedded samples showed an RNA quality of 7.3-8.4 RIN, demonstrating that the RNA quality of worms undergoing re-embedding treatment was not impacted by the re-embedding procedure.

### Spatial Transcriptomics Tissue collection

Methanol fixed, OCT-embedded *Brugia malayi* adult female worm pieces (see section “Parasite material and treatment”) were cryosectioned longitudinally to 8 µm thickness with a CryoStar NX70 cryostat (ThermoFisher). Samples BM1 and BM2 were sectioned with the re-embedding technique described in the section “Total RNA extraction - RNA quality” while sample BM3 was sectioned from the original OCT block. We collected as many sections as possible (30 total) from the 3 worm samples (BM1, BM2, and BM3) onto Visium Spatial Gene Expression Slides (10X Genomics, PN: 2000233). Slides containing tissue sections were stored at −80°C overnight before experimental processing.

### Modified H&E Staining

Spatial Transcriptomics on *B. malayi* adult female reproductive tissue was performed as specified by the 10X Genomics Visium Spatial Gene Expression User Guide^30^, with the following described modifications. Slides with tissue sections were removed from the −80°C, placed on dry ice in a sealed container, incubated at 37°C for 5 minutes in a Thermoblock (ThermMixer with Thermoblock, Eppendorf) with a heated lid (ThermoTop, Eppendorf), and then placed into an ArrayIT metallic hybridization cassette (ArrayIt, AHC1X16) for steps 1.2.d to 1.2.x of the protocol. Step 1.2.b was prepared as stated. For step 1.2.d, 75µL of isopropanol (Millipore Sigma, I9516-25ML) was added to each tissue section well, making sure each tissue section was uniformly covered by the solution. Step 1.2.e was performed as stated. Instead of steps 1.2.f-12.g, isoporopanol (Millipore Sigma, I9516-25ML) was removed from each well and then each section was washed with 75µL of RNase and DNase free MQ water, and the wash was repeated two times for a total of three washes. Step 1.2.h was performed as stated. The slide was then warmed at 37°C for 1.5 minutes in a Thermoblock (ThermMixer with Thermoblock, Eppendorf) with a heated lid (ThermoTop, Eppendorf). For step 1.2.i, 75µL of Mayer’s Hematoxylin (Agilent Technologies, S330930-2) was added to each tissue section well, making sure each tissue section was uniformly covered by the solution. For step 1.2.j, the slide was incubated for 3 minutes at room temperature. Instead of steps 1.2.k-1.2.o, Mayer’s Hematoxylin (Agilent Technologies, S330930-2) was removed from each well and then each section was washed with 75µL of RNase and DNase free MQ water, and the wash was repeated three times for a total of four washes. The slide was then air dried and warmed at 37°C for 1.5 minutes in a Thermoblock (ThermMixer with Thermoblock, Eppendorf) with a heated lid (ThermoTop, Eppendorf). For step 1.2.p, 75µL of Dako Bluing Buffer (Agilent Technologies, CS70230-2) was added to each tissue section well, making sure each tissue section was uniformly covered by the solution. For step 1.2.q, the slide was incubated for 1 minute at room temperature. Instead of steps 1.2.r-1.2.t, the Dako Bluing Buffer (Agilent Technologies, CS70230-2) was removed from each well and then each well washed with 75µL of RNase and DNase free MQ water, and the wash was repeated two times for a total of three washes. The slide was air dried and then warmed at 37°C for 1.5 minutes in a Thermoblock (ThermMixer with Thermoblock, Eppendorf) with a heated lid (ThermoTop, Eppendorf). For step 1.2.u, 75µL of Eosin Mix (100µL Eosin Y Solution + 900µL Tris-Acetic Acid Buffer (0.45 M, pH 6.0)) was added to each tissue section well, making sure each tissue section was uniformly covered by the solution. For step 1.2.v, the slide was incubated for 45 seconds at room temperature. Instead of steps 1.2.w-1.2.y, the Eosin mix (100µL Eosin Y Solution + 900µL Tris-Acetic Acid Buffer (0.45 M, pH 6.0)) was removed from each well and then each well washed with 75µL of RNase and DNase free MQ water, the wash was the repeated two times for a total of three washes. The slide was then removed from the ArrayIT metallic hybridization cassette (ArrayIt, AHC1X16) and air dried. Step 1.2.z was performed as stated. A cover glass (Menzel-Gläser, PN: 12392108, 22×22mm #1, 631-1339) was mounted on the slide with 280µL of 85% glycerol (Sigma, 49767-100ML).

### Microscope & Imaging

For imaging, first an overview 20x image was taken, followed by each individual tissue section imaged with 10 z-stack planes, 2 µm apart at 20x. Hematoxylin & Eosin brightfield images were acquired with a Zeiss Axiolmager.Z2 VSlide Microscope using the Metasystems VSlide scanning system with Metafer 5 v3.14.179 and VSlide software. The microscope had an upright architecture, and used a widefield system; a 20x air objective with the numerical aperture (NA) 0.80 was used. The camera was a CoolCube 4m with a Scientific CMOS (complementary metal-oxide-semiconductor) architecture and monochrome with a 3.45 × 3.45 µm pixel size. All brightfield images were taken with a Camera Gain of 1.0 and an Integration Time/Exposure time of 0.00004-0.00008 seconds.

### cDNA synthesis & Library Construction

After imaging, the 10X Genomics Visium Spatial Gene Expression User Guide^30^ protocol was resumed from step 2.1.a using a Visium Spatial Gene Expression assay (10X Genomics) kit, with the modifications specified here. For step 2.1.e, the tissue sections were permeabilized for 2 minutes. For step 4.2.d, 19 cycles were used to amplify the cDNA for each sample subarray. For step 5.5.d, 14 cycles were used for the Sample Index PCR for BM1 and BM2 samples and 15 cycles for sample BM3.

### Sequencing

The average library length was assessed using the BioAnalyzer DNA High Sensitivity kit (Agilent, 5067-4626) on an Agilent 2100 BioAnalyzer. Library concentration was determined with a Qubit dsDNA BR Assay kit (Thermo Fisher Scientific, Q32850). Libraries were diluted to 2nM, pooled, and sequenced on an Illumina NextSeq 2000 with paired-end, dual index sequencing. Read 1 was sequenced for 28 cycles and Read 2 was sequenced with 150 cycles. Run type and parameters following those specified in the Visium Spatial Gene Expression User Guide sequencing instructions^30^.

## Data Analysis

### Data Pre-processing

TSO adaptor sequences from the 5’ end of transcripts and poly(A) sequences from the 3’ end of transcripts in the Read 2 raw fastq sequence files were trimmed using cutadapt (v2.3) with a custom bash script (https://github.com/ludvigla/VisiumTrim). TSO sequences were defined as a non-internal 5’ adapter with an error tolerance of 0.1, poly-A homopolymers were defined as a regular 3’ adapter of 10 As with an error tolerance of 0, and the minimum overlap was defined as 5 bp. Sequence quality was evaluated before and after trimming with FastQC v0.11.8^40^ and MultiQC v1.8^41^. 10X Genomics Loupe Browser v4.0.0 was used to manually select spots under tissue sections in the H&E jpeg images. Spots under any portion of tissue were selected to maximize the initial number of spots in the dataset. The resulting json files can be found in the Mendeley dataset specified under “Data Availability”.

### Data Processing - Generation of raw counts

Genomic sequence and annotation files for *B. malayi* were acquired from WormBase: WBPS14 and for *Wolbachia* were acquired as RefSeq: NC_006833.1. The *Brugia malayi* genome GFF3 annotation file (brugia_malayi.PRJNA10729.WBPS14.annotations.gff3.gz) was converted to GTF using gffread (with parameters -T -o) from cufflinks (v2.2.1); the GTF file was then filtered to remove all history exons (any exon labeled “history” in the second column of the GTF file). The *Wolbachia* gtf annotation file (GCF 000008385.1 ASM838v1 genomic.gtf.gz) was used to map to *Wolbachia* tRNA and rRNA genes to visualize areas of *Brugia malayi* that contained *Wolbachia*. The *Wolbachia* gtf annotation file (GCF 000008385.1 ASM838v1 genomic.gtf.gz) was modified to change “CDS”‘ in column 3 to “exon” to include all genes in the index. 10X Genomics Space Ranger v1.2.0 was used to build a combined *Brugia malayi*-*Wolbachia* reference with spaceranger mkref and specifying each organism genome fasta file and annotation gtf file as inputs. TSO and poly(A) adaptor trimmed paired fastq files were processed with 10X Genomics Space Ranger v1.2.0 with spaceranger count and inputting the corresponding Hematoxylin & Eosin (H&E) images in jpeg format, the custom *Brugia malayi*-*Wolbachia* reference, and the image manual alignment json file output from 10X Genomics Loupe Browser v4.0.0.

### Quality Control - Data Filtering

The filtered count matrices (filtered_feature_bc_matrix.h5) and tissue H&E images output from Space Ranger v1.2.0 were analyzed in R (v4.0.3) *Wolbachia* and *B. malayi* genes were separated into different count matrices. *B. malayi* gene types were based on gene annotations (these annotation files can be found in the github repository specified under “Code availability”).

Protein coding genes were defined as “protein coding” in the “biotype” column of the annotation file. Mitochondrial and ribosomal protein coding genes were defined by the “annotation” column. The *B. malayi* genes were filtered to contain only protein coding genes, and mitochondrial and ribosomal protein coding genes were filtered out. The *B. malayi* data was further filtered by removing spots with fewer than 30 unique genes, fewer than 50 and greater than 10,000 unique transcript molecules (UMIs), more than 3% of UMIs belonging to mitochondrial genes, and more than 30% of UMIs belonging to ribosomal genes and removing genes with less than 1 spot per gene and less than 1 unique transcript molecule (UMI) count per gene. The *Wolbachia* count matrix was filtered to remove the same spots as were filtered out from the *B. malayi* count matrix. Data filtering was performed using the *STUtility* package (v1.0)^42^ that works on top of the toolkit for single cell analysis Seurat (v3.2.3)^43^ in R (v4.0.3).

### Data Normalization

Each worm sample was split into its own filtered count matrix for normalization. Normalization was performed using the SCTransform function from Seurat with default parameters except unique gene counts regressed out and not returning only variable genes (vars.to.regress = “nFeature_RNA”, return.only.var.genes = FALSE). After normalization, individual sample count matrices were integrated using Seurat functions SelectIntegrationFeatures with default parameters except “nfeatures = 7000” to acquire the integration features, followed by the PrepSCTIntegration Seurat function with default parameters. MergeSTData was run with default parameters to merge the normalized count matrices into a single object. Normalization was performed using the *STUtility* package (v1.0)^42^ that works on top of the toolkit for single cell analysis Seurat (v3.2.3)^43^ in R (v4.0.3).

### Clustering Analysis

On the SCTransform normalized count matrix, dimensionality reduction was performed using Principal Component Analysis (PCA) with RunPCA on the SCT assay and the integration features (output from SelectIntegrationFeatures function in the “Data Normalization” section) as features. Sample batch effects were removed using RunHarmony^44^ (group.by.vars and vars_use as worm_sample) applied on the PCA-computed matrix using 10 dimensions, theta of 0, and maximum iteration of 80. Clustering was performed with Seurat functions FindNeighbors, FindClusters, and RunUMAP on the harmony integrated data with 10 dimensions (dims = 1:10), a k parameter of 10 (k.param = 10), and resolution of 0.3 (resolution = 0.3). Clustering analysis was performed using the *STUtility* package (v1.0)^42^ that works on top of the toolkit for single cell analysis Seurat (v3.2.3)^43^ in R (v4.0.3).

### Differential Expression Analysis

To find the differentially expressed marker genes per cluster, the Seurat function “FindAllMarkers” was used with default settings, except for specifying the SCT assay (assay = “SCT”) and a log fold change threshold of 0.1 (logfc.threshold = 0.1). Only upregulated and downregulated genes with p-value < 0.05 were considered. Differential expression analysis was performed using the *STUtility* package (v1.0)^42^ that works on top of the toolkit for single cell analysis Seurat (v3.2.3)^43^ in R (v4.0.3). We used a previous proteomics study^22^ (the list of proteins enriched in each dissected tissue can be obtained in the WormMine database) to annotate the different clusters based on the differentially expressed (DE) marker genes for each cluster. The previous proteomic study indicated a set of proteins that were enriched in specific tissues^22^, specifically the body wall, digestive tract, and reproductive tract. The proteins enriched for each of these specific tissue type(s) can be considered as markers for those tissues where they were highly expressed. Therefore, we used these markers to validate our clustering analysis. We looked at the proportion of these tissue specific marker genes in each cluster to assign a cluster to a specific tissue type.

### Functional Term Enrichment Analysis

For the functional term enrichment analysis, InterPro description and GO terms for each gene were identified using BioMart^45^. Significantly over-represented functional terms in each cluster were identified using Fisher’s exact test (FDR <0.05).

### Co-localization analysis

We considered spots where at least 1 *Wolbachia* gene UMI was detected as *Wolbachia*+ spots and the spots that did not capture *Wolbachia* were considered *Wolbachia*-negative. Differential Expression Analysis (DEA) was run between *Wolbachia*+ and *Wolbachia*-spots using the Seurat function “FindMarkers” with both wilcox and DESeq2 tests using default settings, except for specifying the SCT assay (assay = “SCT”), a log fold change threshold of 0.1 (logfc.threshold = 0.1), with ident.1 set to *Wolbachia*+ spots and ident.2 to *Wolbachia*-spots. All DESeq2 DE genes were also found by wilcox, and values from both with p-value < 0.05 are included in **Supplementary Table 5**. Colocalization analysis was performed using the *STUtility* package (v1.0)^42^ that works on top of the toolkit for single cell analysis Seurat (v3.2.3)^43^ in R (v4.0.3).

### 3D Figure

The tissue images used in the alignment were pre-processed using Adobe Photoshop 2022 (v23.2.1) to remove background and to separate each section of the worm into one image. The posterior and anterior regions of the separated worm sections were manually aligned using ‘ManualAlignImages()’ and the 3D model was made using ‘Create3DStack()’ from *STUtility* package (v1.0). The 3D images for the genes of interest were generated using ‘FeaturePlot3D()’, also from the *STUtility* package (v1.0) in R (v4.2.0).

### Shiny app

The gene expression heatmap images on the shinyapp were generated using “ST.FeaturePlot()” (dark.theme = T for better color contrast to aid the visualization) and cluster images using “FeatureOverlay()” from the *STUtility* package (v1.0) in R (v4.2.0). Shiny theme ‘sandstone’ was applied as part of the overall design aesthetics. The app is hosted on https://www.shinyapps.io for public access.

### Statistics and reproducibility

The spatial transcriptomics data presented in the Main and Supplementary Figures are generated from *n* = 3 biological replicates of *Brugia malayi* adult female worms.

## Data availability

Raw sample sequence fastq files are available on NCBI SRA under the accession PRJNA870734. Processed gene count matrices, related metadata, corresponding ST tissue H&E microscopy images, and 3D model HTML files are available in the Mendeley dataset under Reserved DOI: 10.17632/8f62vydg3z.1.

The gene expression information for all the *B. malayi* genes from the normalized SCT assay are available for visualization on our publicly available app: https://giacomellolabst.shinyapps.io/brugiast-shiny/

## Code availability

The scripts used to generate count matrices from raw sequence fastq files and related R scripts used to analyze count matrices for quality control and filtering, normalization, clustering and differential expression analysis, colocalization analysis, and 3D figure generation can be accessed from our github repository Brugia_malayi_study (https://github.com/giacomellolab/Brugia_malayi_study).

## Acknowledgements

This work was supported in part by the Division of Intramural Research (DIR) of the NIAID/NIH (D.V., E.G.). S.G. was supported by the Swedish Research Council VR grant 2020-04864. The *Brugia malayi* parasites (NR-48892) were provided by the NIH/NIAID Filariasis Research Reagent Resource Center for distribution through BEI Resources, NIAID, NIH. We thank Uppsala Multidisciplinary Center for Advanced Computational Science (UPPMAX) for providing computational infrastructure. We thank Ludvig Larsson for his discussions on analysis of the Spatial Transcriptomics data.

## Author Information

## Author’s Contributions

H.S. and D.V. performed investigatory experiments, developed methodology, analyzed and visualized the data, worked on validation, curated the data, conducted project administration, and wrote the manuscript. H.S. and M.C. implemented software for formal analysis of the data. Y.M. generated the 3D visualization, set up the shiny app analyses, and gave input for the formal analysis of the data. S.S. aided in methodology and software implementation for formal analysis of the data. S.G. and E.G. conceptualized, designed, led and supervised the study, developed methodology, acquired funding and resources, conducted project administration, and wrote and edited the manuscript.

## Ethics declarations

