## Supplementary Information for "Miniature spatial transcriptomics for studying parasite-endosymbiont relationships at the micro scale"

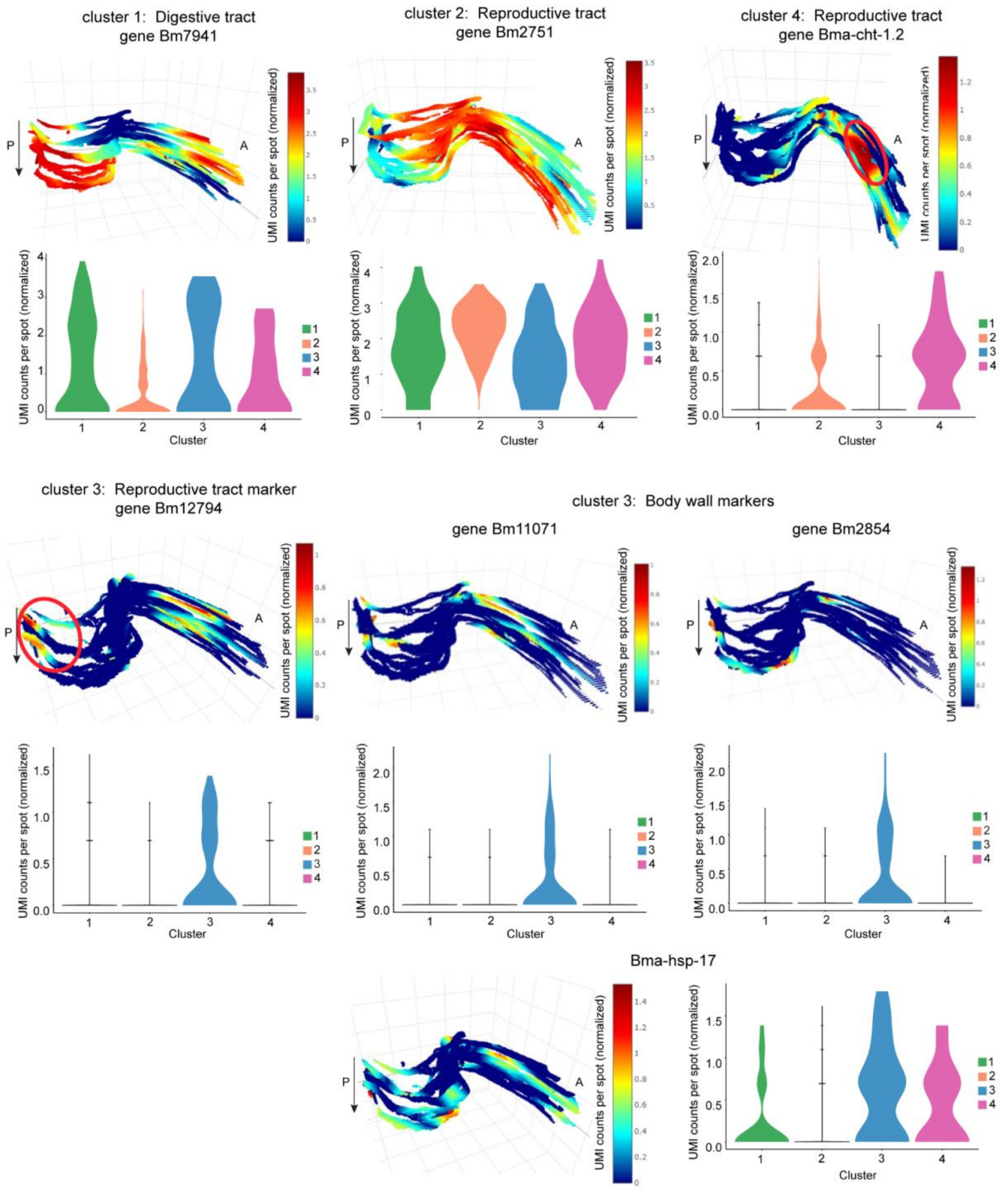


**Supplementary Figure 1. 3D gene expression of cluster marker genes.** Cluster marker genes spatial gene expression in 3D through worm sample BM2 and expression distribution across each cluster in violin plot. Magnification 20x. A: Anterior, P: Posterior, arrow indicates first to last section through the worm.

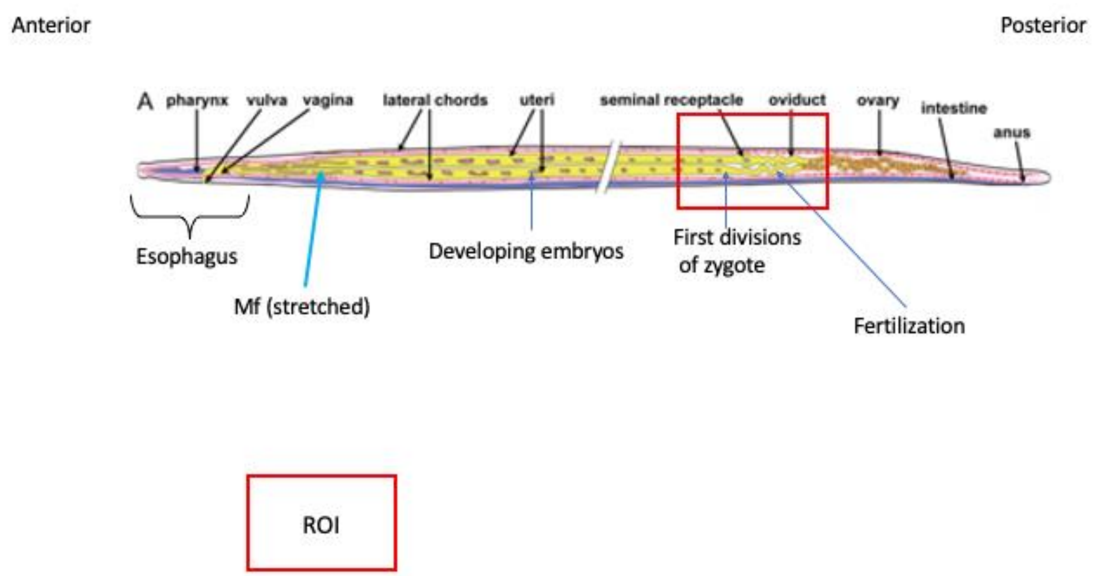


**Supplementary Figure 2. Region of interest (ROI) used in the study.** Schematic of the region of interest used in the study consisting of the posterior region of adult female *Brugia malayi* worms containing ovary tissue, the beginning of the uterus with fertilized eggs and early embryos, digestive tract, and body wall [1].

**Supplementary Table 1**. **Spatial transcriptomics summary.** Raw sequence library information for the samples and sample sections used in the study as output from 10X Genomics Space Ranger.

**Supplementary Table 2. Clustering analysis differentially expressed genes.** Differentially expressed (DE) marker genes for each “miniatureST” cluster. **A.** DE marker genes per cluster. **B.** DE marker genes corresponding to body wall (BW) markers in cluster 3. **C.** DE marker genes corresponding to reproductive tract (RT) markers in cluster 3. **D.** DE marker genes corresponding to digestive tract (DT) markers in cluster 1. **E.** DE marker genes corresponding to reproductive tract (RT) markers in cluster 2. **F.** DE marker genes corresponding to reproductive tract (RT) markers in cluster 4.

**Supplementary Table 3. Fixed term enrichment analysis.** Fixed term enrichment analysis results showing the genes and processes enriched in each cluster. **A.** All results for Fixed term enrichment analysis. **B.** Fixed term enrichment analysis results included in Figure 2E.

**Supplementary Table 4. Pathways of interest genes.** Genes associated with glycolysis, gluconeogenesis, lactate dehydrogenase, and enzymes that convert cysteine amino acids to pyruvate.

**Supplementary Table 5. Colocalization analysis differentially expressed genes.** Differentially expressed (DE) genes in *Wolbachia*+ versus *Wolbachia*- spots.
